# Metabolic costs of walking and arm reaching in persons with mild multiple sclerosis

**DOI:** 10.1101/2022.08.31.506034

**Authors:** Robert Courter, Enrique Alvarez, Roger Enoka, Alaa Ahmed

## Abstract

Movement slowness is a common and disruptive symptom of multiple sclerosis (MS). A potential cause is that individuals with MS slow down to conserve energy as a behavioral adjustment to heightened metabolic costs of movement. To investigate this prospect, we measured the metabolic costs of both walking and seated arm reaching at five speeds in persons with mild MS (pwMS; n = 13; 46.0 ± 7.7yrs) and sex- and age-matched controls (HCs; n = 13; 45.8 ± 7.8yrs). Notably, the cohort of pwMS was highly mobile and no individuals required a cane or aid when walking. We found that the net metabolic power of walking was approximately 20% higher for pwMS across all speeds (p = 0.0185). In contrast, we found no differences in the gross costs of reaching between pwMS and HCs (p = 0.492). Collectively, our results suggest that abnormal slowness of movement in MS – particularly reaching – is not the consequence of heightened effort costs alone. Our findings are consistent with the possibility that demyelination of reward regions of the central nervous system in MS disrupt the dopamine-mediated impetus to move more quickly and thereby prompt slower movements.

**NEW & NOTEWORTHY:** Individuals with multiple sclerosis (MS) often move more slowly than those without the disease. A possible cause is that movements in MS are more energetically expensive and slowing is an adaptation to conserve metabolic resources. Here, we find that while walking is more costly for persons with MS, arm reaching movements are not. These results bring into question the driving force of movement slowness in MS and implicate other motor-related networks contributing to slowing.

## Introduction

An inflammatory disease of the central nervous system (CNS), multiple sclerosis (MS) elicits a myriad of sensory, cognitive, and motor symptoms. The symptoms are consequences of axonal demyelination, or potential total neurodegeneration, which disrupts efferent and afferent signal transmission between brain and body (1–3). One of the primary barriers to our understanding of MS rests in its heterogeneous presentation of neurological disability dependent upon the location of these demyelinated lesions within the CNS (4).

Despite this high person-to-person variability, a prevalent symptom experienced by persons with MS (pwMS) is movement slowness in walking (5–9), upper limb movements (10–14), and even saccades (15–17). There are, however, several potential causes of movement slowness in pwMS that likely depend on the location of inflammation in the CNS. Slowness in walking or reaching could be due to pyramidal tract lesions leading to spasticity (18) or due to motor unit loss contributing to muscle force unsteadiness (19, 20); whereas saccade dysfunction could be linked to cerebellar damage causing tremor or dysmetria (17). Rather than pursue these specific possibilities, we instead approach it from a higher-level, neuroeconomics perspective.

Both the selection of a movement and the subsequent speed with which it is performed is akin to an economic transaction: spending effort to purchase a more rewarding bodily state (21). The fundamental goal of movement is then to maximize the net reward rate (22), such that we should move more quickly to obtain a reward despite incurring greater effort costs (23). In healthy adults and nonhuman primates, for example, an expectation of reward produces dopamine-mediated faster reaction times and quicker movements in both reaching (21, 24, 25) and saccades (26–28). Conversely, added effort in the form of metabolic cost leads to slower walking and reaching, while preferred speeds are often selected to minimize the associated metabolic effort (29–35). Hence, movements can be successfully modeled in terms of their abstract cost function – maximizing reward intake while minimizing effort expenditure – which can assist in understanding CNS organization of motor behavior in the healthy brain, or when neurological disease disrupts this control.

The possibility we investigated was that pwMS move more slowly in walking and reaching to conserve metabolic energy and maintain a higher net reward rate. The energetic costs of walking (5, 7, 9, 36–39) and other mobility tasks (40, 41) trend higher for pwMS, thus it is feasible that walking slowness in MS could be credited to higher effort. However, it is often difficult to pinpoint the sources of these elevated metabolic costs.

One possibility for increased walking effort in pwMS is lower fitness or diminished exercise tolerance. The resting metabolic rate for pwMS does not appear to be higher than healthy adults (37, 39, 42), suggesting that disparities in energy expenditure may be a result of reduced physiological fitness due to more sedentary lifestyles (i.e., a secondary consequence of other symptoms, such as fatigue or pain) (9, 42–46). The 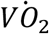 max or peak, a marker of cardiorespiratory fitness, in arm-crank or cycle ergometry tasks for pwMS tends to be lower than healthy controls, while pwMS tend to rely on a greater proportion of their 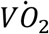 capacity to perform a task (9, 41, 42 44, 46). Distinguishing between any direct pathophysiological changes of MS against indirect reductions in fitness on the metabolic costs of movement remains unknown.

A movement that is less metabolically costly than walking is seated arm reaching (32, 47), but arm reaching energetics have never been explored for pwMS despite large percentages of individuals reporting upper limb impairment (48). Accordingly, the purpose of our study was to investigate whether the metabolic costs of both walking and seated arm reaching are higher for persons with mild MS as compared to healthy age- and sex-matched control participants (HCs). If so, this would suggest that movement slowing in pwMS in both walking and reaching is a rational response to increased metabolic costs.

We used indirect calorimetry and systematically measured the metabolic costs of walking and reaching across five different speeds. Critically, our participants were rigorously screened to ensure that individuals could walk at speeds up to 1.6 m/s without the use of a cane or other walking aid. Our primary hypotheses were two-fold: *(i)* the metabolic costs of walking would be higher for pwMS; and *(ii)* the costs of seated reaching would not differ for pwMS and HCs due to the low cardiorespiratory demand of the task.

## Methods

### Participants

Participants with MS (pwMS) (n = 13; 46.0 ± 7.7 yrs; 11F) and healthy, age- and sex-matched (±3 years) controls (HCs) (n = 13; 45.8 ± 7.8 yrs; 11F) were recruited upon passing a strict pre-screening process and participated after informed consent was obtained. All procedures were approved by the University of Colorado Boulder Institutional Review Board (Protocol #20-0072).

Primary inclusion criteria involved: *(a)* between the ages of 18-65 years old; *(b)* within ±3 years of age to a pwMS (HC-group only); *(c)* clinical diagnosis of MS (pwMS-group only) and free of other neurological disorders; *(d)* PDDS score ≤ 3 or ambulatory without an assistive device (pwMS-group only); and *(e)* on stable doses of symptomatic-treating medications (pwMS-group only). Primary exclusion criteria involved: *(a)* a relapse or administration of systemic steroids within the last 30 days (pwMS-group only); or *(b)* presence of other risk factors, including other major diseases, cognitive impairment, mental disorders, drug use, spasticity, or recent orthopedic injury that would otherwise limit exercise involvement.

### Protocol

Participants visited the lab on two 2 separate occasions with 7 to 14 days between visits. On the first day of testing, participants fasted 2-4 hours in advance. After obtaining written, informed consent, participants first completed a battery of questionnaires (MFIS, PROMIS Sleep Disturbance, PROMIS Sleep Impairment, and MSWS-12) and physical assessments (weight, GPT, MVC grip, and T25FW).

A standing baseline metabolic trial was performed which involved 5 minutes of quiet standing on the treadmill while breathing through the metabolic mouthpiece. Participants then walked on the treadmill at five speeds (0.6, 0.85, 1.1, 1.35, and 1.6 m/s) for five minutes each while breathing into the mouthpiece (Figure 1A). The order of speeds was randomized, and participants were required to take at least 5 min of seated rest between walking conditions. While walking, participants were instructed to “breathe and walk as naturally as possible” and asked to refrain from holding the handrails of the treadmill, if possible. Participants performed a second, standing baseline metabolic trial at the conclusion of the first visit.

**Figure 1.**
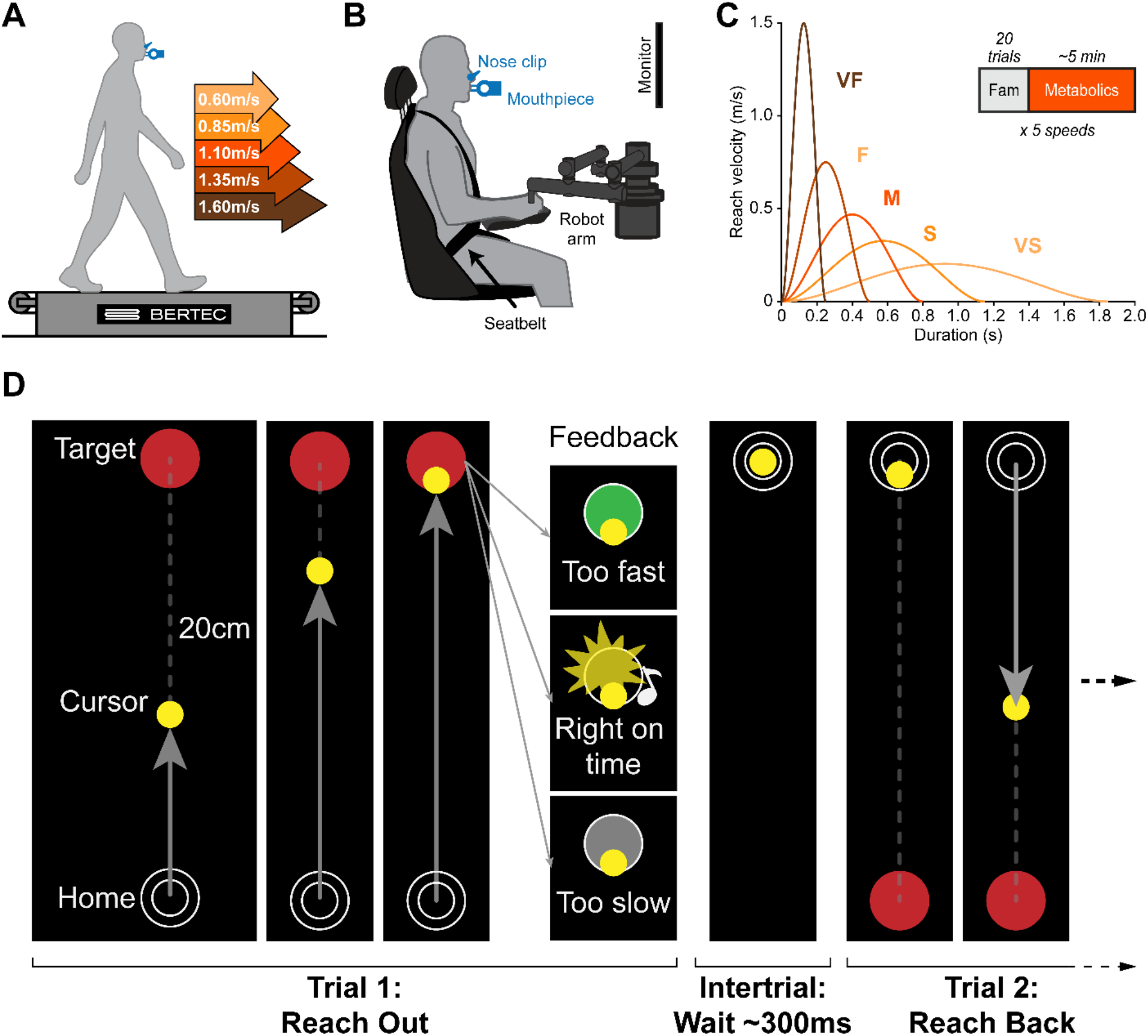
Experimental methods. Metabolics were collected via indirect calorimetry while participants: **A.** Walked on the treadmill at five speeds, ranging from 0.6 to 1.6m/s; and **B.** performed horizontal planar arm reaching movements at five speeds while grasping the handle of the robot arm. **C.** Simulated minimum jerk velocity trajectories for a 20cm reach completed within the five enforced time durations. The enforced times to complete a reach in each speed condition were 1850, 1150, 800, 500, and 250ms ± 50ms corresponding to the “very slow” (VS), “slow” (S), “medium” (M), “fast” (F), and “very fast” (VF) conditions, respectively. **D.** Audiovisual display presented on the monitor during reaching. Participants controlled the yellow cursor with the robot arm and moved from the home circle to the target circle. Upon arrival at the target circle, participants received differing audiovisual feedback dependent on whether they arrived at the target “Too fast,” “Right on time,” or “Too slow.” After a 300ms intertrial interval in which participants kept the cursor still in the target circle, a new target appeared in the previous home location and the next trial began.

The second day of testing involved reaching metabolics. Participants were required to fast for a longer period (6-8 hours) beforehand, typically overnight. Participants then made seated reaching movements using their dominant arm while breathing into the mouthpiece (Figure 1B). When reaching, participants grasped the handle of the shoulder-elbow robot and controlled the position of a virtual cursor displayed on a vertically positioned monitor at eye level. Reaches were 20 cm in distance and performed in an out-and-back fashion along an anteroposterior axis originating near the participant’s sternum. Like with walking, participants reached at five different speeds ranging from “very slow” (~10 cm/s) to “very fast” (~80 cm/s) for at least five minutes each (Figure 1C). The order of speeds was again randomized, and participants were required to take at least 5 min of seated rest between conditions. Participants completed a seated baseline metabolic trial at the beginning and at the conclusion of the visit.

Unlike walking on the treadmill, participants could not be forced to reach a given speed. To enforce reaching speeds, we used audiovisual feedback to encourage participants to reach from one target to the next in a specified amount of time. The position of the handle controlled a cursor (r = 0.3 cm) on a vertical computer monitor in front of the subject positioned at eye level. To begin a trial, participants moved the cursor within the home circle (r = 0.8 cm) and remained still for 300 ms. After this 300 ms delay, the home circle disappeared, and a red circular target (r = 0.8 cm) appeared 20 cm away. Participants were then instructed to reach “out” (i.e., primarily elbow extension) to the new target with this cue and stop within it. After another 300 ms delay with the cursor inside the target, the next target appeared in the position of the previous home circle, and the reach “back” (i.e., primarily elbow flexion) could begin. Audiovisual feedback was provided when the cursor reached the target and was based on the time it took for the participant to move the 20 cm from the home circle to the target circle. There were five duration requirements which consequently enforced average reaching speed: 1850, 1150, 800, 500, and 250 ms corresponding to the “very slow,” “slow,” “medium,” “fast,” and “very fast” conditions, respectively (Figure 1C). If the cursor reached the target within ± 50 ms of the specified duration, the target would flash yellow and play a pleasant tone. If the cursor reached the target outside of this ± 50 ms time window, the target would flash gray or green, indicating the movement was “too slow” or “too fast,” respectively (Figure 1D). Participants learned to associate the audiovisual stimuli to the speed of their reaching movements during a 50-trial practice block prior to beginning the main reaching protocol. Each reaching condition also began with 20 familiarization trials followed by a 30 s break to allow participants to experience the upcoming speed.

### Data Acquisition

#### Metabolic cost

Gas exchange (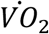 and *VĊO*_2_) was measured breath-by-breath by the metabolic cart (TrueOne2400, ParvoMedics, Sandy, UT, USA). Participants wore a nose clip and breathed through a 2-way, nonrebreathable value (Hans Rudolph, Kansas City, MO, USA) while walking on a split-belt treadmill (Bertec, Columbus, OH, USA) and reaching with the arm while grasping the handle of a shoulder-elbow robot (Interactive Motion Technologies, Watertown, MA, USA) (Figure 1A, B). Participants were required to fast before metabolic testing (2-4 hours prior to walking; 6-8 hours or overnight prior to reaching) and performed any given walking or reaching movements for a minimum of five minutes to allow metabolic steady state to be reached. The metabolic cart 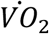 and *VĊO*_2_ sensors were calibrated before each visit using certified gas mixtures, and the flowmeter calibrated using flow rates from a 3L syringe. Baseline metabolic rates were measured for a minimum of five minutes while quietly standing and quietly sitting both at the start and end of each of the walking and reaching visits, respectively.

The 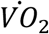 and *VĊO*_2_ exchange values over the final two minutes of each walking or reaching condition were averaged. Gross metabolic rates (W) were then estimated using the averaged steady state gas exchange values in the Brockway equation (49). Net metabolic rates (W) were estimated by subtracting the lower of the two baseline metabolic rates from the gross metabolic rates. Normalized metabolic rates were calculated by dividing the gross or net rates by the participant’s body mass (W/kg).

#### Reaching Kinematics

Robot handle position and velocity data was acquired at 200 Hz. Position data was filtered using a fourth-order Butterworth filter with 10 Hz cutoff. Velocity, acceleration, and jerk along each axis were obtained by differentiating the filtered position signals. Movement onset was determined using a custom algorithm that used a threshold on the standard deviation of tangential velocity. Movement offset was determined as the first timepoint that the cursor traveled 20 cm along the anteroposterior axis.

#### Fatigue

The self-reported level of fatigue was measured with the 21-item Modified Fatigue Impact Scale (MFIS) (50–52). Questions are answered based on how fatigue had impacted an aspect of day-to-day life ranging from 0 (*Never*) to 4 (*Almost always*) within the past four weeks. We used the total MFIS score, which incorporates the physical, cognitive, and psychosocial subscales. Higher scores indicate increased feelings of fatigue, and a cutoff score of 38 is often used to discriminate “fatigued” from “non-fatigued” individuals (51).

#### Sleep Quality

Aspects of sleep quality and its impact were measured with two questionnaires: the Patient-Reported Outcomes Measurement Information System (PROMIS) Sleep-Related Impairment and Sleep Disturbance scales (53, 54). The PROMIS Sleep-Related Impairment is a 16-item self-assessment measuring the impact of sleep on quality of life in the past seven days and is rated on a five-point Likert scale ranging from “*Not at all”* to *“Very much.”* Similarly, the PROMIS Sleep Disturbance is a 27-item self-assessment measuring the quality of sleep within the past seven days and is rated on the same aforementioned five-point Likert scale. Higher scores indicated worse sleep and heightened detriments owing to poor sleep. Sleep quality was measured due to its high correlation with fatigue (55).

#### Walking ability

Self-assessed walking ability was measured with two questionnaires: the Patient Determined Disease Steps (PDDS) (56) and MS Walking Scale (MSWS-12) (57). The PDDS is a 1-item assessment asking how well pwMS can walk ranging from 0 (normal) to 8 (bedridden). We used a PDDS score of ≆ 3 (*walking only mildly impaired, do not typically require a cane/walking aid*) as a primary inclusion criterion. The MSWS-12 is a 12-item assessment asking pwMS about their limitations with walking in the past two weeks and is rated on a five-point Likert scale ranging from 1 (*Not at all*) to 5 (*Extremely*).

Maximum walking speed was measured with the Timed 25-foot Walk Test (T25FW) (58). The time it took participants to walk a prescribed 25-foot distance “as quickly, but as safely, as possible” was timed with a stopwatch. The fastest of two trials was recorded.

#### Upper limb ability

Upper-limb ability was measured with two physical assessments: the Grooved Pegboard test (GPT) (59, 60) and maximal voluntary contraction (MVC) grip (61). The GPT provided a measure of manual dexterity. Participants placed 25 pegs into the holes as quickly as possible, twice with each hand in alternating fashion. The fastest time for each hand was recorded. The MVC grip provided a measure of hand-grip strength. Participants held a dynamometer and squeezed as hard as possible, three times with each hand in alternating fashion. The largest MVC (kg) for each hand was recorded.

### Statistical analyses

Data were analyzed using R version 4.0.5 (62). All HCs were able to complete all reaching and walking speeds. All pwMS were able to complete the five reaching speeds and slowest three walking speeds; 12 of 13 pwMS were able to complete the second fastest (1.35 m/s) and 11 of 13 the fastest (1.6 m/s) walking speeds. One pwMS did not complete the second day of reach testing. These uncompleted speeds were treated as missing data points in subsequent statistical analyses.

Participant demographics, questionnaire scores, and baseline physical assessment performance are described in Table 1. Numeric outcome variables were compared with the non-parametric 2-sample Kolmogorov-Smirnov test. Nominal data were compared with the χ^2^-test. Effect sizes were calculated with Cohen’s *d*. Comparisons of reach kinematics (i.e., speed, accuracy, and jerk) were made using two-sample t-tests and p-values were reported with Bonferroni corrections for multiple comparisons.

**Table 1.** Participant characteristics.

| Variable* | HC<br>(N = 13) | MS<br>(N = 13) | p-value | Effect Size<br>(95% CI) |
| --- | --- | --- | --- | --- |
| *Physiological Assessments* |  |  |  |  |
| Sex |  |  |  |  |
| F | 11 (85%) | 11 (85%) | -- | -- |
| M | 2 (15%) | 2 (15%) | -- | -- |
| Age (years) | 45.9 (7.8)<sup>†</sup> | 46 (7.7) | 1<sup>‡</sup> | -0.02 (-0.83; 0.79) |
| Height (cm) | 166.6 (6.2) | 166.1 (8.7) | 0.88 | 0.06 (-0.75; 0.87) |
| Mass (kg) | 65.9 (7.6) | 76.9 (17.9) | 0.13 | -0.80 (-1.64; 0.04) |
| MS Diagnosis (years) | -- | 9.37 (8.52) | -- | -- |
| MVC Grip (kg) | 34.2 (7.5) | 32.1 (7.8) | 0.57 | 0.27 (-0.54; 1.09) |
| Pegboard Time (s) | 54.7 (7.6) | 61.9 (10.0) | 0.13 | -0.80 (-1.64; 0.04) |
| T25FW (s) | 3.42 (0.45) | 4.33 (1.13) | **0.015** | -1.05 (-1.91; -0.19) |
| *Questionnaires* |  |  |  |  |
| MFIS | 9.6 (14.8) | 36.7 (20.6) | **0.0039** | -1.51 (-2.43; -0.59) |
| PDDS | -- | 1.23 (1.17) | -- | -- |
| MSWS-12 | -- | 28 (11.40) | -- | -- |
| Sleep Disturbance | 49.2 (18.2) | 64.7 (24.0) | 0.13 | -0.73 (-1.56; 0.11) |
| Sleep Impairment | 26 (10.7) | 40.2 (18.7) | 0.13 | -0.93 (-1.78; -0.08) |

\*Abbreviations - HC: Healthy controls; MS: Multiple sclerosis; MVC Grip: Maximum voluntary contraction grip force; T25FW: Timed 25-foot Walk; MFIS: Modified Fatigue Impact Scale; PDDS: Patient Determined Disease Steps; MSWS-12: MS Walking Scale. <sup>†</sup>Values reported as mean (SD). <sup>‡</sup>p-values and effect sizes from nonparametric 2-sample Kolmogorov-Smirnov test and Cohen's *d*, respectively.

Walking and reaching metabolic data were analyzed using linear mixed effects regression models (LMER) predicting metabolic power. Due to the established nonlinear relationship between movement speed and metabolic power in both walking and reaching (33, 47), power was log-transformed to best satisfy the standard regression assumptions of linearity, homoscedasticity, and normality. Fixed effects were the movement speed, a binary MS indicator (pwMS = 1), and their interaction. Random intercepts on participant were included to capture within-subject variability. Interactions were kept in the models regardless of significance, due to the models’ purpose being testing of *a priori* hypotheses:

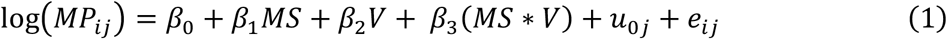

LMERs with both log-transformed power and log-transformed walking speed, as well as gamma-distributed generalized linear mixed effects (GLMER) models with log-links were also tested and compared to our LMERs; however, the LMERs were deemed to provide better overall fits via residual distributions, log-likelihoods, and AIC/BIC scores, though results did not significantly change across choice of model.

Exploratory statistical analyses were performed to investigate the potential modulatory effect of other factors (i.e., demographic data, questionnaire scores, physical ability, etc.) on the relationship between metabolic power, speed, and MS. We used R’s built-in *step* function (62) to perform a backwards elimination procedure to build stepwise LMER models predicting walking or reaching metabolic power from a specified full model containing all possible fixed predictor variables and random effects.

## Results

### Walking energetics were greater in PwMS

The log-transformed net power normalized to body mass (W/kg) increased with walking speed (*β_speed_* = 1.039; p < 2e-16) (Figure 2B). On average, a 0.1 m/s increase in walking speed resulted in a 10.95% (95% CI: [10.26, 11.65]) increase in net normalized power. A significant main effect of group suggested that pwMS used an average of 20.54% (95% CI: [3.33, 40.60]) more metabolic power for walking at a given speed than HCs (*β_MS_* = 0.187; p = 0.0185) (Figure 2B). The group-by-speed interaction (*β*_*MS:Speed*_ = −0.084; p = 0.0709) suggested that differences in net normalized power between pwMS and HCs started to converge at faster walking speeds, though was nonsignificant.

**Figure 2.**
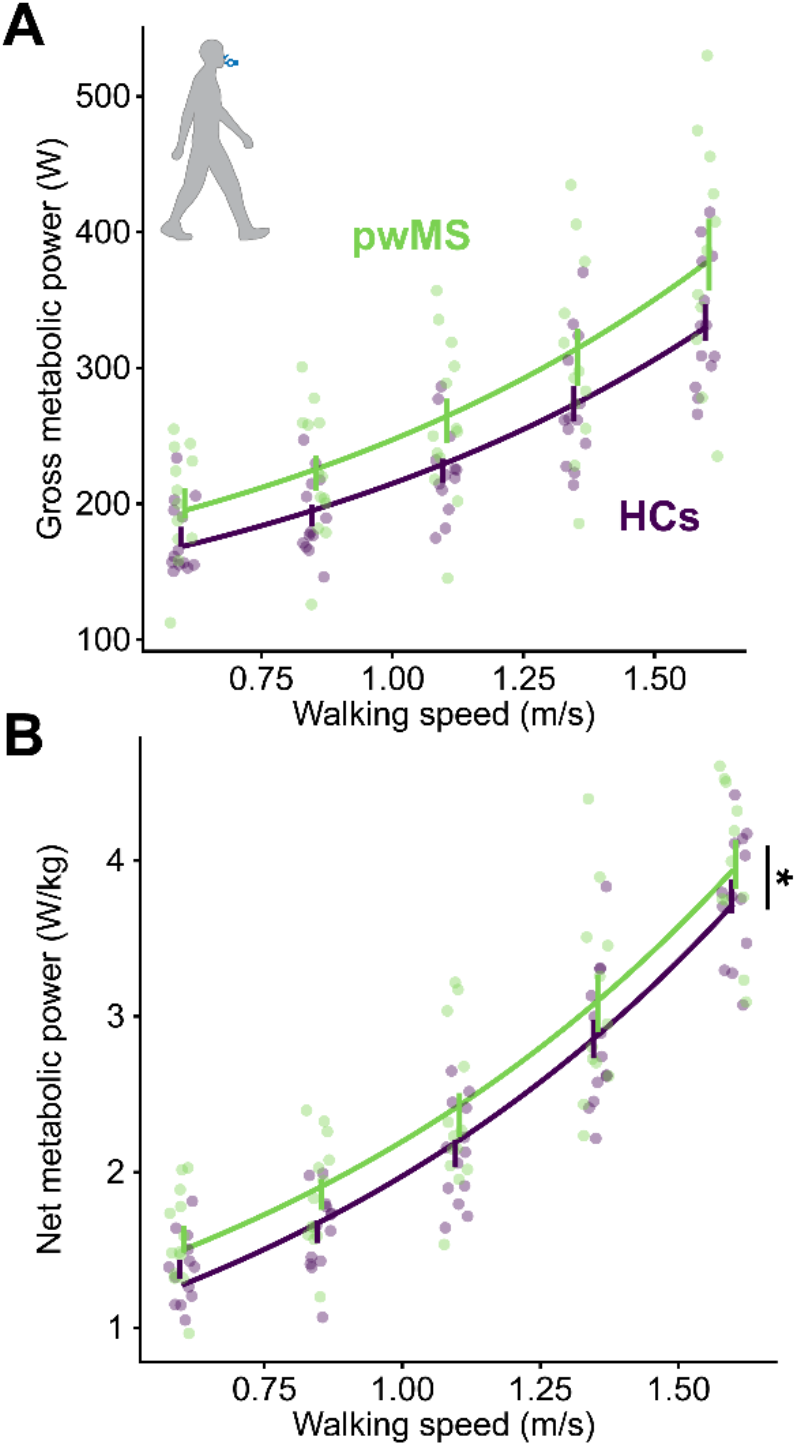
Walking metabolic power. **A.** Gross metabolic power of walking increased non-linearly with speed. Participants with MS (pwMS; green) required more gross metabolic power to walk at a given speed, on average, as compared to healthy controls (HCs; purple) **B.** Same as in **A** but displaying net metabolic power of walking normalized to body mass. (*Effect of group is significant at the 0.05 level).

Net metabolic power normalized to body mass is a standard metric for walking. Nonetheless, we also analyzed the gross costs to account for any changes in baseline rates that may have occurred over the course of the protocol. Log-transformed gross power (W) also increased linearly with speed (*β_Speed_* = 0.656; p < 2e-16). A nonsignificant effect of group suggested that pwMS require around 14.00% (95% CI: [-2.78, 33.66]) more gross metabolic power for walking at a given speed than HCs, on average (*β_MS_* = 0.131; p = 0.106) (Figure 2A).

### Reaching energetics are similar in HC and PwMS

The log-transformed gross power (W) increased with reaching speed (*β_Speed_* = 1.798; p < 2e-16). On average, a 0.1 m/s increase in hand speed resulted in a 19.70% (95% CI [17.60, 21.83]) increase in gross power (Figure 3A). There was no main effect of group (*β_MS_* = −0.053; p = 0.492), which suggests that pwMS and HCs had similar average costs of reaching at a given speed. For example, the −5.21% (95% CI [-18.77, 10.65]) less average metabolic power to reach at a given speed for pwMS was not significantly different than the HCs (Figure 3A). Similar to walking, the group-by-speed interaction suggested that metabolic power increased more steeply with speed for pwMS, though again was nonsignificant (*β_MS:Speed_* = 0.207; p = 0.147).

**Figure 3.**
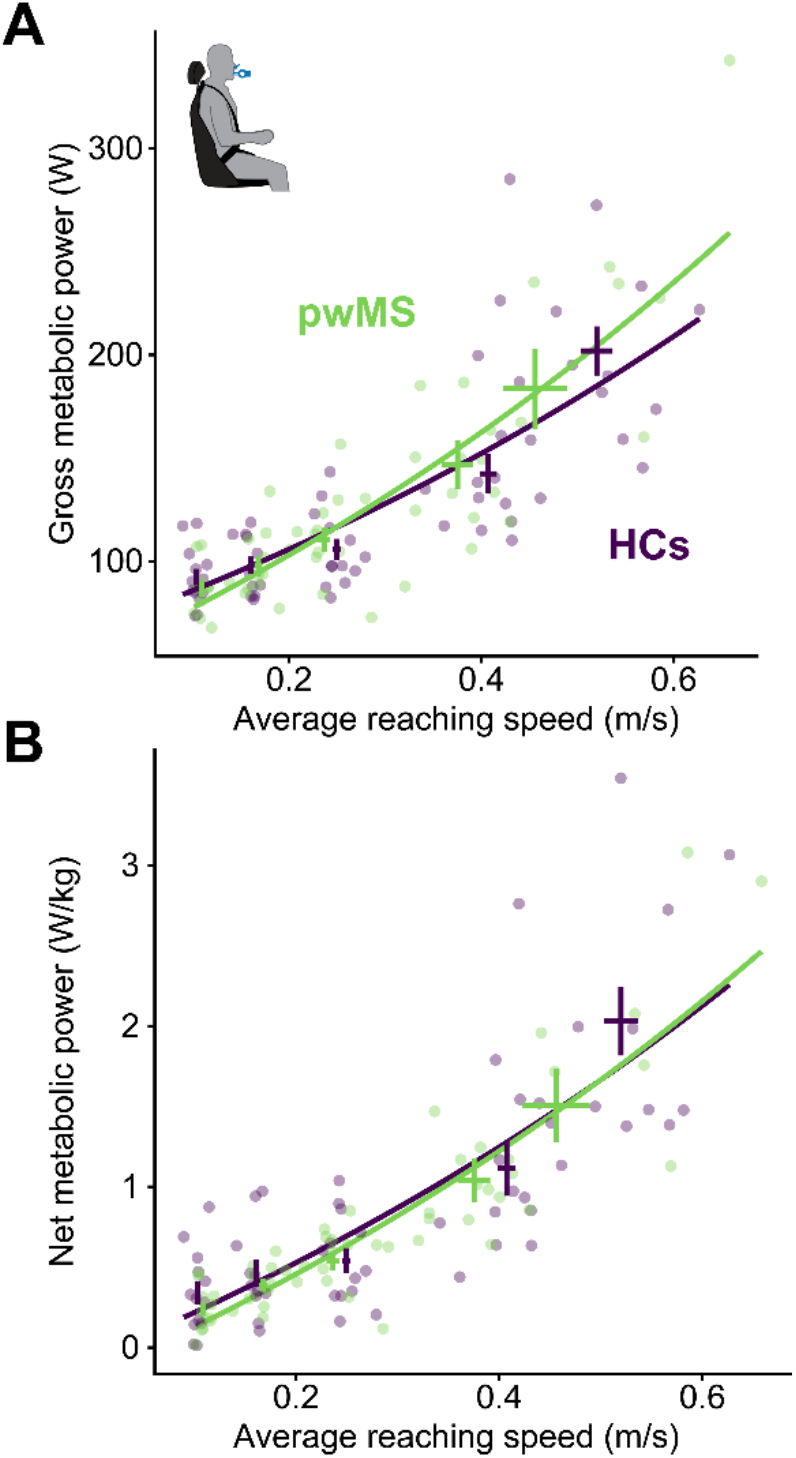
Reaching metabolic power. **A.** Gross metabolic power of reaching increased non-linearly with speed. Participants with MS (pwMS; green) and healthy controls (HCs; purple) required similar average gross power to reach at a given speed. Because reach speed could not be enforced as exactly as walking speed, crosses indicate the average power (± s.e.) at the average reaching speed (± s.e.) within a condition. **B.** Same as in **A** but displaying net metabolic power of reaching normalized to body mass.

The metabolic costs of reaching are reported as gross metabolic power, because, in contrast to walking, mass does not need to be transported during reaching tasks and net costs are modest. Nonetheless, we did analyze the net metabolic power of reaching normalized to mass as is standard in locomotion. Log-transformed net normalized metabolic power (W/kg) of reaching increased linearly with reaching speed (*β_Speed_* = 4.91; p < 2e-16). The effect of group remained unchanged (*β_MS_* = −0.033; p = 0.898), suggesting that pwMS and HCs had similar net costs of reaching at a given speed (Figure 3B).

We compared reaching kinematics between pwMS and HCs across the five speed conditions (“very slow” to “very fast”) (Figure 4A, B). There were no significant differences in average reaching duration (Figure 4C) or speed (Figure 4D) between pwMS and HCs in any of the five conditions (p > 0.05). We also compared reach performance and kinematics in each speed condition. There were no significant differences in average sum of squared jerk between pwMS and HCs for any reach speed (p > 0.05) (Figure 4F). Reach accuracy was calculated as the Euclidean distance from the cursor to the center of the target at the first moment velocity went below 2.5 cm/s. Reach accuracy did not significantly differ between pwMS and HCs in any condition (Figure 4E). Together, the similar average speeds, jerks, and accuracies suggested that any findings in reaching metabolics were likely not credited to kinematic differences. Furthermore, use of LMERs in lieu of repeated measures ANOVAs enabled us to use participant’s actual average speed per condition, rather than binning into an ordinal “speed condition” variable.

**Figure 4.**
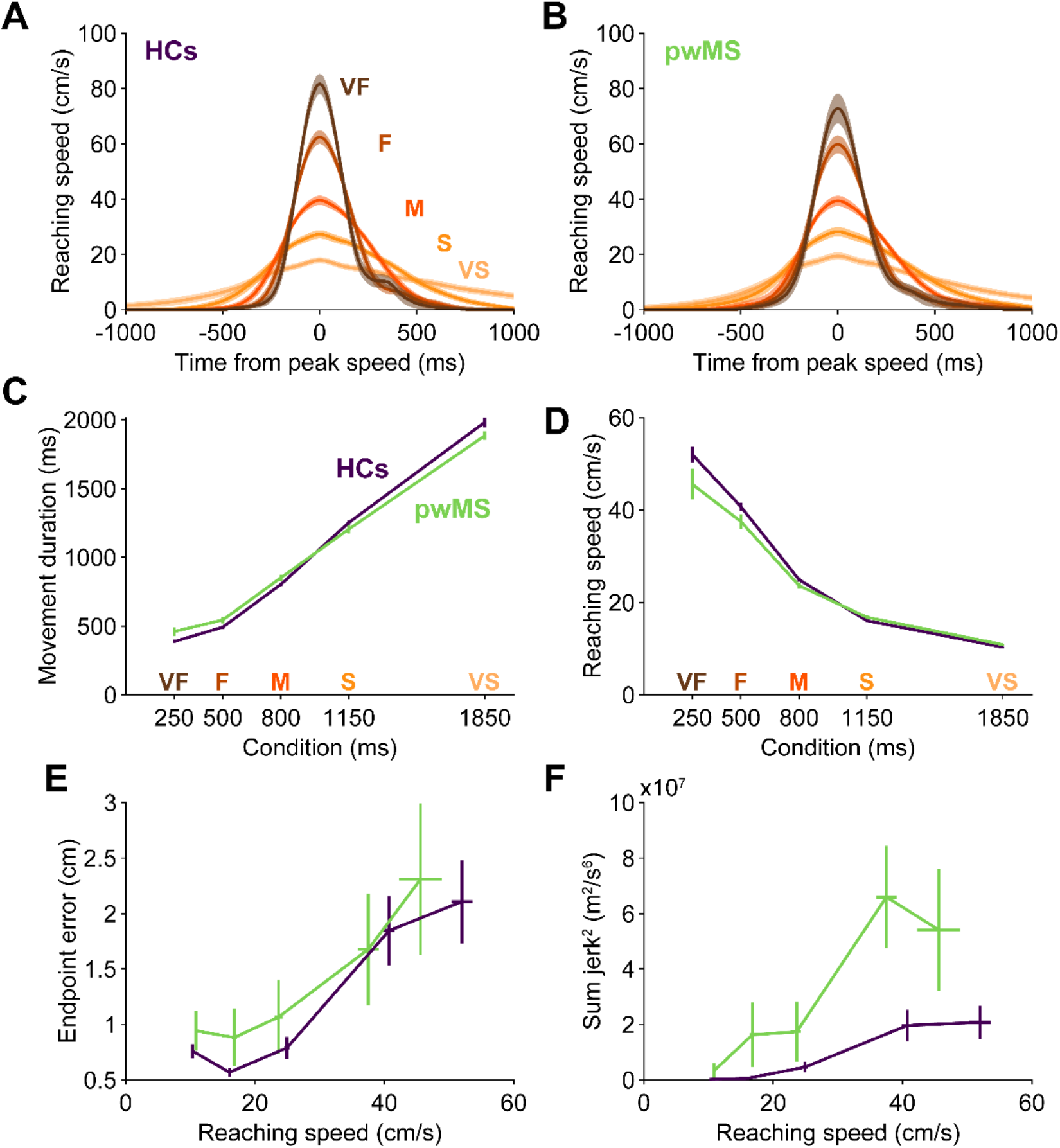
Reaching kinematics. **A-B.** Average reaching speed profiles in the “very slow” (VS), “slow” (S), “medium” (M), “fast” (F), and “very fast” (VF) conditions in **(A)** healthy controls (HCs) and **(B)** participants with MS (pwMS) aligned to peak speed. **C-D.** Average **(C)** movement durations and **(D)** reaching speeds did not significantly differ between HCs (purple) and pwMS (green). **E.** Average endpoint error in each condition did not differ between HCs (purple) and pwMS (green). The endpoint of a reach was defined as the last moment reach speed exceed 2.5cm/s. **F.** Average sum of squared jerk did not significantly differ between HCs (purple) and pwMS (green). Shading **(A & B)**, vertical **(C-F)**, and horizontal (**E-F)** error bars represent mean ± s.e.

### Baseline metabolic rates are similar between HC and PwMS

Previous work has suggested that the baseline 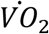 or metabolic rates do not differ between pwMS and HCs (37, 39, 42). We found that the standing (MS: 79.72 ± 13.36W; HC: 84.39 ± 10.67 W) and sitting (MS: 68.73 ± 9.98W; HC: 69.93 ± 10.32W) baseline rates were not different between groups (p > 0.05). However, the baseline rate while standing was significantly higher than that of sitting for both HCs and pwMS (p < 0.05). Note that the average baselines in the sitting and standing positions were quantitatively less for pwMS than HCs – this slight discrepancy possibly helps explain why the net normalized metabolic power of walking was significantly higher for pwMS (*β_MS_* = 0.187; p = 0.0185), but gross metabolic power was not (*β_MS_* = 0.131; p = 0.106). This further implicates reduced exercise tolerance or fitness contributing to elevated walking energetics in pwMS.

### Participant characteristics

Comparisons between the 13 pwMS and 13 HCs are enumerated in Table 1. Briefly, neither age, height, nor weight differed between groups (p > 0.05). There were no differences in PROMIS Sleep-Related Impairment or Sleep Disturbance scores between pwMS and HCs. Neither grooved pegboard times nor MVC grip of the dominant hand differed between groups (p > 0.05). Self-reported fatigue measured via the MFIS was higher for pwMS (MS: 36.69 ± 20.62; HC: 9.62 ± 14.79; p = 0.0039). Seven of thirteen pwMS and one of thirteen HCs scored higher than the standard cutoff score of 38, indicating a clinically relevant level of fatigue (51, 63).

Time since initial disease diagnosis for the pwMS-group was 9.37 ± 8.52yrs. PwMS scored 1.23 ± 1.17 on the PDDS and 28 ± 11.40 on the MSWS-12, indicating overall low levels of mobility impairment. Also, T25FW time was still slower for pwMS (MS: 4.33 ± 1.13s; HC: 3.42 ± 0.45s; p = 0.0022).

### Exploratory analyses

To account for the potential modulatory influence of participant characteristics and covariates on metabolic costs, we performed stepwise regression via backwards elimination to construct LMERs predicting metabolic costs from a full set of potential predictor variables.

Only four predictor variables were selected from a full set of fourteen for the walking analysis (Figure 5A). Log-transformed net power of walking normalized to body mass was predicted to increase with age (*β_Age_* = 0.009; p = 0.0188), increase with MVC grip force of the dominant hand (*β_MVC_* = 0.010; p = 0.0095), and increase with speed (*β_Speed_* = 0.998; p < 2e-16) (Figure 5B). Critically there was a significant main effect of group, again indicating that pwMS required more gross power, on average, than HCs to walk (*β_MS_* = 0.114; p = 0.0368) (Figure 5B):

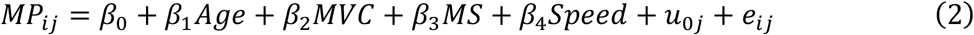

PwMS required 12.07% (95% CI [1.82, 23.35]) more metabolic power than HCs when controlling for potential covariates in this more detailed model. The model explained the variability in the data well, with a marginal *R*^2^ (i.e., considering fixed-effects only) of 0.852 and a conditional *R*^2^ (i.e., considering both fixed and random effects) of 0.946 (64).

**Figure 5.**
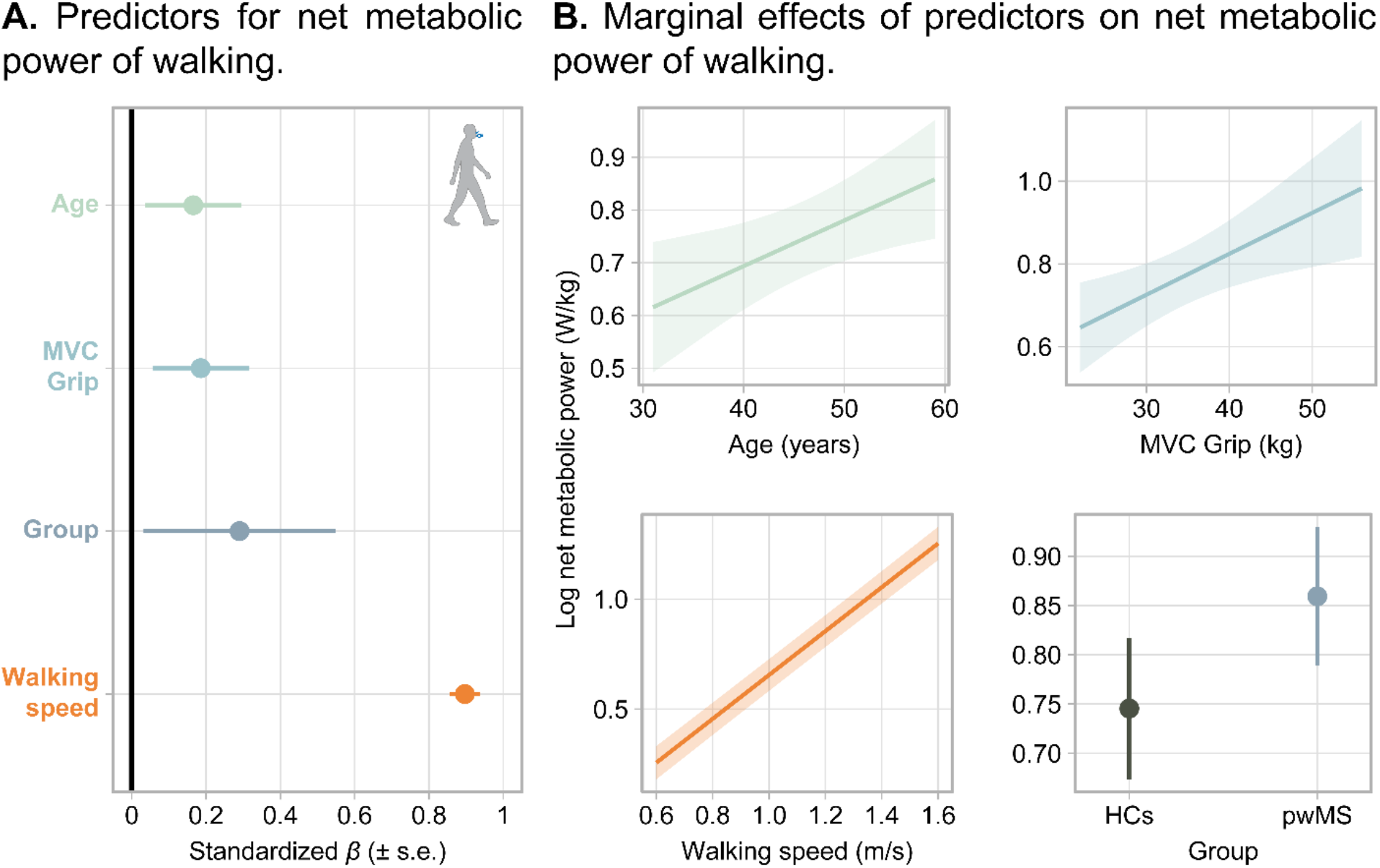
Stepwise LMER model predicting the net metabolic power of walking normalized to body mass. **A.** Standardized coefficients (± s.e.) for the four predictor variables remaining in the final model. A positive value indicates an increase in the predictor results in an increase in net metabolic power. A negative value indicates an increase in the predictor results in a decrease in net power. **B.** Marginal effects (i.e., effect of an individual predictor on the outcome while fixing all other predictor variables) of the four selected predictors. Shaded bands and error bars are 95% confidence intervals for predicted power.

Only three predictor variables were selected out of sixteen for the reaching analysis (Figure 6A). Log-transformed gross metabolic power of reaching was predicted to decrease marginally with an increase in PROMIS Sleep Disturbance score (*β_Sleep Disturb_* = −0.003; p = 0.0475), increase with slower T25FW time (*β_T25FW_* = 0.084; p = 0.0382), and increase with speed (*β_Speed_* = 1.887; p < 2e-16) (Figure 6B):

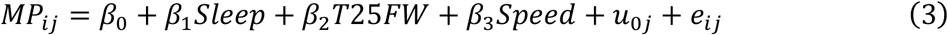

Noticeably, any categorical main effect of MS was discarded from this model, reinforcing the primary finding that pwMS do not exhibit heightened costs of reaching as compared to HCs.

**Figure 6.**
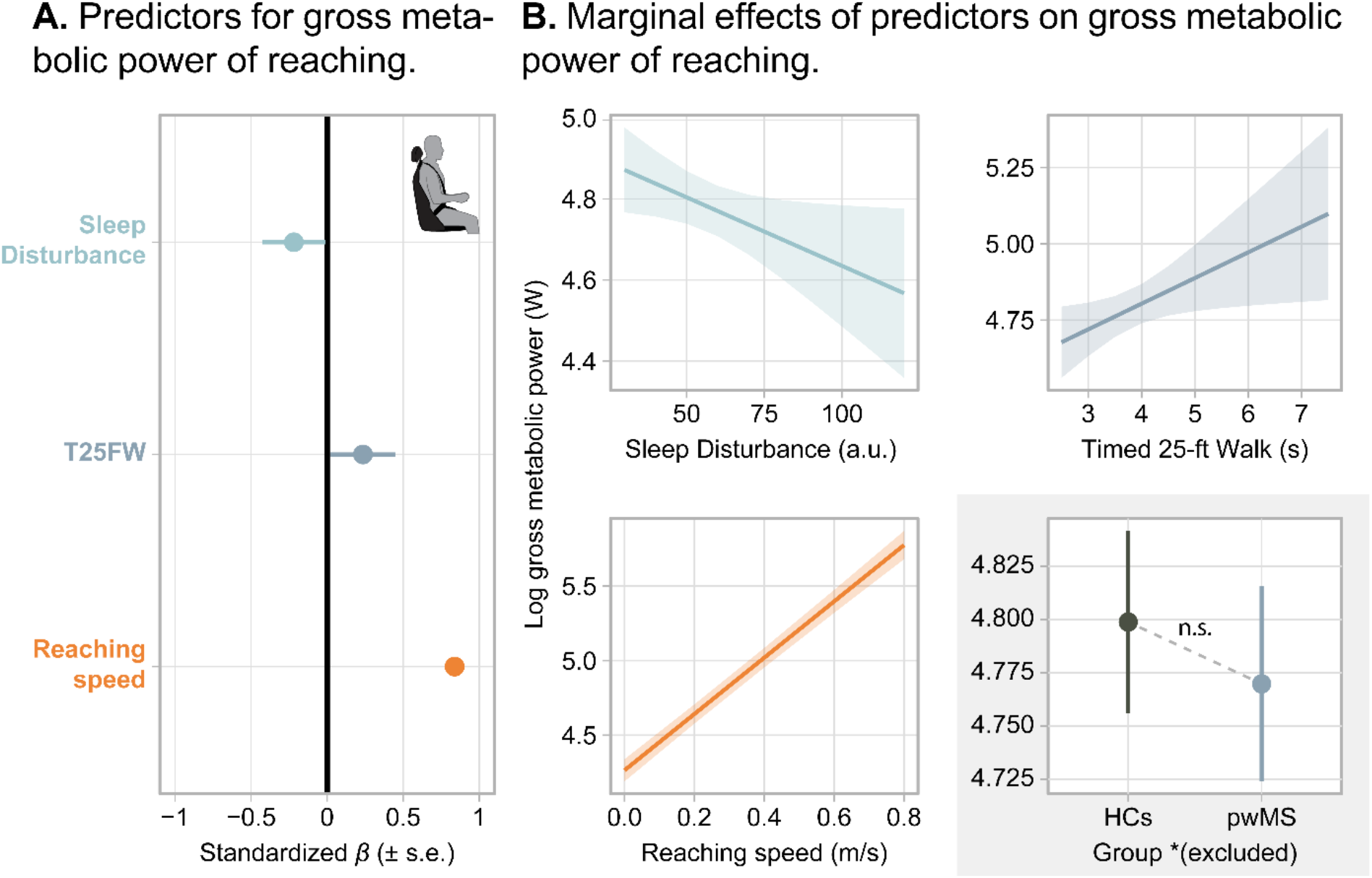
Stepwise LMER model predicting the gross metabolic power of reaching. **A.** Standardized coefficients (± s.e.) for the three predictor variables remaining in the final model. A positive value indicates an increase in the predictor results in an increase in gross metabolic power. A negative value indicates an increase in the predictor results in a decrease in gross power. **B.** Marginal effects (i.e., effect of an individual predictor on the outcome while fixing all other predictor variables) of the three predictors. Shaded bands are 95% confidence intervals for predicted power. *Note that the effect of group (± s.e.) (bottom right) was excluded from the final model and was displayed for sake of comparison.

This model likewise explained the variability in the data well, with a marginal *R*^2^ of 0.714 and a conditional *R*^2^ of 0.894 (64).

## Discussion

Does increased metabolic cost contribute to movement slowing in multiple sclerosis? To answer this question, we measured the metabolic costs of both walking and seated arm reaching movements in persons with mild multiple sclerosis (pwMS) compared to healthy age- and sex-matched controls (HCs). We found that while the metabolic costs of walking were higher at any speed for pwMS, the metabolic demands of seated arm reaching movements were no different than HCs. These results suggest that complementary mechanisms beyond metabolic cost, such as perceived effort or reward valuation, are slowing movements for pwMS.

### Walking costs were elevated for persons with MS

The metabolic cost of walking is contingent on such variables as speed, incline, mass, and age (29, 33, 65 66). We found that the metabolic cost of walking increases with speed at a similar rate in both HCs and pwMS, but that there is an MS-dependent inflation in cost of around 20%.

Our findings diverged slightly from previous studies investigating walking costs in MS. For one, we only found an approximate 20% increase in walking costs for pwMS compared to HCs, which is not as dramatic as previously reported two-to-three fold increases in walking costs (7, 36, 41). This disparity is likely due in-part to the novelty of our dataset: it consists of a relatively homogenous cohort of pwMS who did not require a cane or walking aid, and a majority of whom did not rely on any contact with the handrails of the treadmill. Larger differences in cost were found in studies with thirty (37) to fifty (7) to 100 percent (41) of their MS groups requiring walking aids.

Furthermore, most studies investigating walking in MS use the oxygen cost of walking as their proxy for energy expenditure (5, 7, 9, 36, 37, 39, 41). The oxygen cost of walking is the amount of oxygen required to move one kilogram of mass one meter. Instead, we employed the Brockway equation (49) to convert oxygen and carbon dioxide exchange rates into metabolic power, which may have contributed to our smaller differences in cost.

Work from Chung and colleagues found that the oxygen cost of walking for pwMS was approximately 78.5% higher at a slow walking speed of 0.6 m/s, but not at a preferred (around 1.0 m/s) or fast walking speed of 1.4 m/s (37). While we did not find a convergence of metabolic power at faster speeds, we did see an insignificant interaction term in the walking regression trending in that direction (p = 0.0709). The convergence of costs at faster speeds could be attributed to lower muscular demand required to maintain postural control or lateral stability when compared to slow speeds (37). PwMS experience proprioceptive-based deficits and reduced dynamic stability while walking (67, 68), which correlate negatively with walking speed(9) and encourage pwMS to walk more quickly to exploit the passive dynamics of walking (69).

Comparable increases in walking cost are observed in healthy older adults. Energetic costs of walking for older adults are ~23% higher than young adults (66). For example, young and older adults walking at 1.3 m/s were found to utilize 2.57 ± 0.09 W/kg and 2.96 ± 0.11 W/kg, respectively (70). Our cohort of pwMS, though much younger in age, required ~20% more metabolic energy on average, and 3.08 ± 0.18 W/kg when walking at a similar 1.35 m/s. Corroborating this similarity with older adults, age was selected as a significant predictor of walking metabolic power in the stepwise regression model, such that older individuals expend more energy even when controlling for MS status. MS, although not an age-related disease, heightens locomotor energy demand in a manner mirroring healthy aging.

### Reaching costs were not elevated for persons with MS

Similar to walking, reaching metabolic costs vary with factors such as speed, mass, distance, and age (47, 71). We appear to be the first to measure the metabolic costs of arm reaching in pwMS and demonstrate that across a range of feasible reach speeds, the costs were not related to MS disease state.

The closest analog to reaching metabolics performed previously is arm crank ergometry (44), although this was done in a maximal exercise performance context. When performing arm crank ergometry, pwMS had reduced peak aerobic capacity, weakened respiratory muscles, and impaired cardiopulmonary responses. In arm reaching, despite the absence of a statistically significant difference between groups, we found the gross metabolic power for pwMS appeared to demonstrate a small divergence at the fastest reach speeds: beginning to cost more than HCs at these higher velocities. The body’s cardiopulmonary and metabolic response to physical activity may have come into play at these fastest speeds which, with peak velocities exceeding 1.0 m/s, were approaching the limit of achievable hand speeds. Interestingly, the gross power of reaching at the fastest speed was no different than walking at the slowest in both groups, implicating the physical demand of reaching very quickly is akin to walking slowly. While reaching faster than what we required in the fastest condition would be challenging, pwMS may have greater costs if moving even more quickly; however, for standard day-to-day reaching requirements, the energetic demands during arm reaching do not differ between pwMS and HCs.

Nonetheless, pwMS still tend to reach more slowly. While we did not find significant differences in the jerk or accuracy of reaches between pwMS and HCs, we did see pwMS reaching with quantitatively slower average speeds in the fastest conditions (Fig. 4). A recent study had pwMS and controls perform horizontal planar reaches and found that pwMS reached with slower, jerkier, and more inaccurate movements than controls, while also adopting altered muscle activation patterns and synergies (12). Pellegrino and colleagues recruited a cohort of pwMS with a higher disability status and required participants to reach to one of eight radial targets as accurately as possible (12). We may have not found differences in reach kinematics due to the lower disability of our cohort and the relative simplicity of our task: reaching out-and-back between two targets with endpoint accuracy, though enforced, not as critical to the protocol as was speed-matching accuracy. It remains possible then that pwMS slow upper limb movements to maintain task accuracy in the face of compromised sensorimotor upper limb control (72, 73).

### Pathophysiological changes in metabolism caused by MS

Why are walking but not reaching costs elevated in MS? One explanation is that the costs of walking for pwMS are heightened due to worsened physiological fitness. Deconditioning occurs as a secondary consequence of the symptoms of MS, such as fatigue, which disincentives exertion and leads to a more sedentary lifestyle. Physiological fitness is often quantified in domains such as aerobic capacity, balance, and muscle strength, all of which tend to be negatively impacted by MS(9, 44, 46) and all of which may increase walking demands. Exercise-based rehabilitation has been shown to both reduce the energetic cost of walking and increase the preferred speeds in pwMS (45), and more active pwMS have lower costs than those less active (38). However, whether the benefits of exercise restore locomotory performance to levels comparable to control individuals is uncertain. Regardless of the exact mechanism, the slower gait speeds in pwMS could be explained in-part by the increase in metabolic effort as speed is reduced to minimize costs (36, 74).

A second explanation is that there are distinct metabolic shifts caused directly by MS pathology. There can be irreversible neurological damage that occurs in MS which causes permanent mobility disability (75), or compensatory axonal mechanisms that alter central nervous system (CNS) energetic demand (76). Thus, despite adequate rehabilitation, pwMS may neither regain full mobility nor restore typical energetic costs of movement due to axonal injury, loss of motor units, decayed muscle strength, and other such adaptations (19, 20).

We measured the metabolic costs of walking and arm reaching with the justification that seated arm reaching movements are far less costly and may remove a large proportion of the fitness-based changes in energetics. The lack of difference in reaching costs suggests that these MS-related pathophysiological shifts in CNS metabolism were, at the very least, not detectable, if occurring at all, and that worsened attributes of fitness may be a principal basis of expensive walking.

### Implication of reward-related CNS regions contributing to movement slowing

If the goal of movement is to maximize a reward-effort tradeoff over time (i.e., a net reward rate) (21), then upper limb slowness (10–14) in MS is not accounted for through metabolic effort. One possibility is that instead individuals are appraising effort on a relative scale. Similar to the fitness hypothesis for walking differences, pwMS have a lower aerobic capacity (42, 44, 46), thus equivalent gross metabolic costs of reaching in pwMS and HCs would equate to a larger proportion of maximal capability for pwMS. Perhaps, then, pwMS are reaching and walking more slowly as a behavioral response to diminished physiological potential.

Alternatively there is reward, which increases movement speed so that we may maximize the rate of reward accrual (25–28). Mechanistically, dopamine tends to encode reward or its expectation (77–79) in regions of the basal ganglia. Dopamine is released moments before movement onset increases the excitability of primary motor cortex (80, 81), facilitating more vigorous movements.

Recent evidence suggests that gray matter demyelination and impaired dopamine mediation within the basal ganglia may emerge as the disease progresses (2, 82–86). PwMS present smaller cortical gray matter volume, and deep gray matter atrophy – consisting of thalamus, putamen, globus pallidus, caudate, and amygdala – tracks with disease progression (84, 85). Accordingly, pwMS improperly value reward and make non-optimal reward-based decisions (87–91). Diminished reward responsiveness in pwMS may hinder assessment of expected reward rate and slowing movement because of an undervalued reward-effort tradeoff. As a result, pwMS may move more slowly due to a paucity of reward-mediated motivation; however, this link between reward, dopamine, and movement invigoration in MS remains an open question.

### Limitations

Our approach is not without its limitations. One shortcoming is that we did not collect self-reported measures of perceived effort or state fatigue (50) between blocks of walking or reaching. Using a Visual Analog Scale (VAS) or Borg RPE would have provided us with a measure of how subjective task difficulty changed over time and how it compared between pwMS and HCs. Higher perceived effort could be inflating the total effort of movement for pwMS, thus further contributing to movement slowness. In healthy individuals, for example, measures of perceived fatigability tend to correlate with the perceived effort of reaching movements (92), while pwMS tend to subjectively rate exercise tasks as more effortful (41, 93). Despite no differences in physical reaching effort between pwMS and HCs, the possibility exists that reaching is perceived as more effortful in pwMS and contributes to movement slowness.

While not the primary focus of the study, a second limitation is that we did not measure preferred walking or reaching speeds. Both preferred walking and reaching speeds are often selected to minimize the cost of transport (J kg^−1^ m^−1^) of the movement (33, 47, 71). Recent evidence suggests that pwMS minimize walking cost of transport when selecting preferred speeds (45, 74), but that the costs tend to be more expensive than controls. Measuring preferred walking and reaching speeds in this study would have also revealed whether our cohort of pwMS were selecting slower speeds than our HCs. That being said, obtaining preferred speeds, especially in reaching, is not a trivial task (94–96).

Lastly, our cohort of pwMS was a convenience sample of predominantly women (12 of 13) with RRMS, which is not necessarily representative of the overall sex distribution of the disease, which sees females 2-3 times more likely to develop MS (97). Further, the inclusion criteria and advertising description of this fairly strenuous study likely introduced selection-bias in attracting more physically active pwMS (38). However, this same selection-bias may not have been the case in our control group in which walking is likely a more trivial task than for pwMS. Nonetheless, this limitation may serve to strengthen some of our results – increased walking costs were still detected even in a reasonably fit and mobile cohort of pwMS. Recruiting a more physiologically diverse group of participants may have lent itself to even greater differences in metabolic costs of movement. The strict inclusion criteria for this study were largely established to ensure safe walking ability on the treadmill; future work could flexibly expand inclusion criteria especially if only the seated reaching movements are of interest.

## Conclusion

Individuals with mild MS expended approximately 20% more metabolic energy when walking compared to persons without MS, but there was no difference between the two groups in metabolic energy consumption during seated arm reaching. These increased costs of walking were found even in a highly mobile cohort of individuals with MS who did not require walking aids. Moreover, the lack of differences found in reaching costs were not attributed to disease-related altered kinematics: speeds, accuracies, and jerk were comparable between the MS and control groups. Our results suggest that movement slowness occurring with MS is not altogether a consequence of energy conservation. Instead, other MS-related mechanisms, such as magnified perceived effort or diminished reward sensitivity, may additionally compel slower movements and should be considered when treating mobility symptoms.

## Disclosures

The authors declare that they have no conflicts of interest.

## Grants

This work was supported by grants from the National Institutes of Health (1R01NS096083) and the National Science Foundation (CAREER award 1352632)

## Notes

FUNDING: This work was supported by grants from the National Institutes of Health (1R01NS096083) and the National Science Foundation (CAREER award 1352632)

### Competing Interest Statement

The authors have declared no competing interest.

